# ATACofthesnake: A scalable framework for analyzing multifactorial and time course chromatin accessibility data

**DOI:** 10.64898/2026.09.18.752590

**Authors:** Ward Deboutte, Aurelie Lenaerts, Adrian Salatino, Thomas Manke

## Abstract

Advances in sample extraction techniques, improvement of library preparation methods and the dramatic reduction in sequencing costs have made complex, multifactorial experimental design the de facto standard in epigenomic studies. In contrast to the plethora of tools, workflows and libraries available to analyze gene expression data, consolidated workflows for chromatin accessibility data are lacking and analysis of such data often occurs on an ad hoc basis. Here, a scalable end-to-end workflow, ATACofthesnake, is presented that simplifies analysis of multifactorial chromatin accessibility data considerably and provides a newly created framework to handle time series data reproducibly.

## Background

Chromatin accessibility is a crucial determinant in enabling eukaryotic gene transcription and temporally regulated cell fates (1). Its dynamic regulation allows for the organism to coordinate developmental processes (2,3), maintain cellular homeostasis (4) and respond to changes in the environment, such as infections (5), stress (6), tissue damage (7) and aging (8). Given the central role chromatin accessibility plays in the epigenetic regulation of the eukaryotic cell, substantial effort went into the development of high-throughput methods to capture the chromatin accessibility state of cells and tissues, both at the bulk level and with single-cell resolution (9). A breakthrough in this domain was achieved with the creation of the Assay for Transposase Accessible Chromatin with high-throughput sequencing (ATAC-seq) protocol, which drastically decreased the hands-on time and difficulty of sample preparation when compared to previously developed assays (10). To date, the ATAC-seq protocol is considered the gold standard to profile chromatin accessibility in cells and tissues genome wide (11), and numerous adaptations were created to allow accessibility profiling in experiments with low cell numbers (12), fixed cells (13), or complex tissues (14). Together with the reduction of sequencing costs, the threshold to design complex experiments containing multiple treatment arms, genotypes, and batch effect covariates have become the norm rather than being an exception (3,15,16). Even though a number of computational tools were developed to address specific hypotheses such as transcription factor occupancy (17) and nucleosome positioning (18), a general framework that fully exploits chromatin accessibility data and that handles a multifactorial design in a statistically appropriate manner is still lacking. In fact, most complex chromatin accessibility analyses resolve to either simple set differences to determine differential accessible regions between conditions or rely on simple group-wise comparisons that do not take all available information into account. The outcome of both approaches is not ideal, in the sense that crucial statistical information is either simply ignored or omitted. This is exacerbated by the fact that most biological experiments are performed with a small number of replicates and thus have limited statistical power to begin with. Secondly, tools that allow two-group comparisons often make it difficult, if not impossible, to incorporate additional factors inherent to the experimental design, such as nuisance variables, or non-factorial covariates such as time points, ordinal covariates or continuous covariates. Thirdly, having to combine different ad hoc analyses hampers reproducibility and increases the risk for inconsistencies and errors during analyses. To tackle these issues, an end-to-end workflow, ATACofthesnake (AOS), was created that allows researchers to conduct quality control, calculate differential accessible regions between groups taking the full experimental design into account, and model time series either in a continuous fashion or by encoding time points as an ordinal covariate. The entire workflow is written in snakemake (19), making both the execution process as FAIR as possible and documenting analyses trivial (20). Substantial effort went into making the formulation of designs and requesting differential analyses straight forward. More specifically, users request analyses on a covariate level rather than on a model coefficient level, since the translation to a linear contrast happens internally. This omits the need for a user to think in terms of the modeling framework, making the workflow suited for both experimentalists with limited computational and statistical experience, as well as more experienced bioinformaticians.

## Results

### ATACofthesnake provides an end-to-end workflow starting from deduplicated BAM or CRAM files

Given the sheer amount of tools and workflows available to process raw and align FASTQ files (19,21,22), BAM/CRAM files were chosen to be the starting point of ATACofthesnake allowing end-users to adhere to the alignment method, alignment score filtering method and deduplication method of their liking. AOS comprises two computational modules (Figure 1). One of these modules is always executed: functions that perform basic quality control on the data, and the generation of a count matrix based on the union of all called peaks across samples, annotated using gene models that the end-user provides. To facilitate standardized quality assessment, the QC output is formatted to be compatible with MultiQC(23), allowing users to aggregate metrics across large cohorts effortlessly. While the number of required input parameters has been reduced to a minimum to enhance ease of use, both modules are highly customisable via various optional input arguments upon runtime (Supplemental table S1). The second module, the hallmark of AOS, calculates differentially accessible peaks and is triggered by providing both a samplesheet and comparison file. Three different differential modes are possible. The first is a two-way comparison of two groups, which can be defined by specifying them using the factor levels defined in the provided samplesheet. This enables simple two way comparisons, but also allows the end-user to define more complex groups in cases of a multifactorial experimental set up. A second differential mode allows determining differential peaks by executing a likelihood-ratio test (LRT). These can be requested by specifying both a full, and a reduced model where the cofactor(s) of interest are omitted. Both the two-group mode and the LRT mode are executed using the quasi-likelihood estimation framework implemented in edgeR (24). The final mode allows modeling of time course data by utilizing Gaussian process regression. Time course data can be treated either as a continuous covariate or as an ordinal covariate. A permutation module is set up to identify peaks that are time-dependent, or to test differences between (groups of) different covariate levels (see methods). If sufficient differential peaks are found in any of the requested modes, downstream motif analysis and footprinting analysis is automatically performed given that a motif file is provided. Since AOS is written on top of Snakemake, parallel execution becomes very easy to handle. As a result, resources can be readily managed and both runtimes and memory utilization are comparable to other genomic workflows, with the longest process (the permutation analysis in the Gaussian process regression) having a runtime just shy of two hours on a 32-core compute node (64 threads, 3GHz) (Supplemental figure S2).

**Figure 1.**
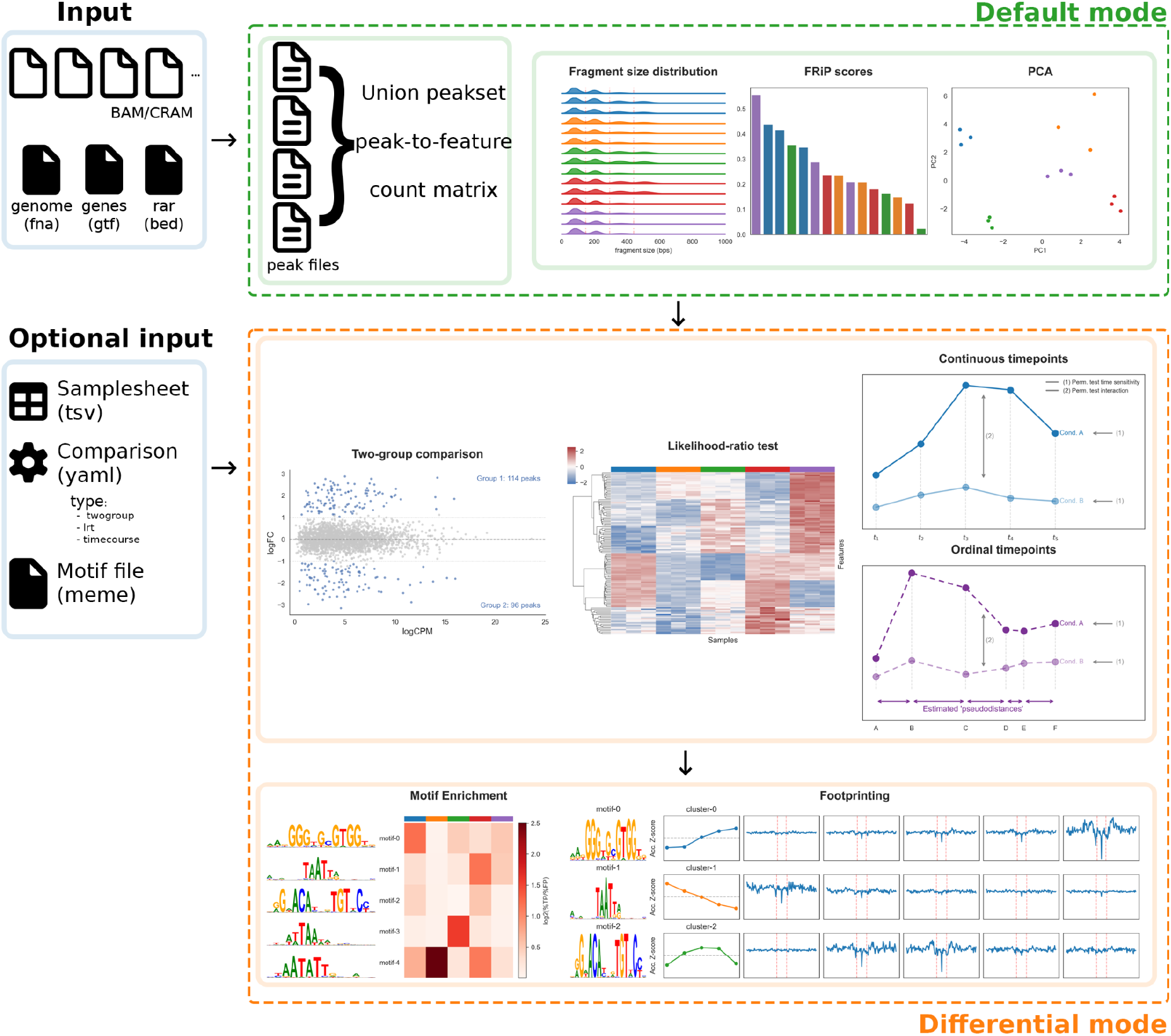
Overview of the AOS workflow. Input file types (both required and optional) are denoted in the blue boxes. The file formats for both required and optional inputs are denoted between brackets. The read-attracting region file is denoted as ‘rar’. The default mode (green) consists of the generation of peak-to-feature annotation files, a unionized peakset across all samples, and a count matrix that serves as input to calculate the QC metrics (denoted as figures in the default mode). The differential mode is denoted in orange and shows the three different modalities (two-group, LRT and time course analyses) as well as the automated downstream motif enrichment and footprinting analysis. Which differential mode or modes are ran can be fully controlled by using the optional input yaml file.

### ATACofthesnake facilitates calling and analysis of differential peaks in a multifactorial setting

To illustrate the ease of use of AOS, public data from the ImmGen project was used to perform both a two-group comparison and a LRT comparison (25). This particular dataset comprises ATAC-seq samples derived from different stages in murine thymopoiesis, a pre-T stage (DN1, DN2, DN3) and a T stage (DP T cells, CD4 T cells and CD8 T cells). In this particular setting there is thus only one covariate (cell type) with six levels. AOS makes it trivial to perform, for example, a pre-T versus T stage comparison, which would otherwise require a formulation of a linear contrast. Requesting this analysis can be done by defining both groups and their corresponding factor levels in the comparison file (Supplemental table S3). In general, two-group comparisons are always specified on their factor level, rather than on the coefficient level. Additionally, the use of the LRT mode allows determining cell type specific peaks without assuming any directionality, by testing against a reduced model containing an intercept only (Supplemental table S3). Both analyses, including the quality control, can be run with a single command. The QC module reveals that the fragment size distribution of all samples show, to some degree, the typical ATAC-seq nucleosomal periodicity (Figure 2A). The fraction of reads falling in peaks (FRiP scores) range between 0.087 and 0.55 (Figure 2B) and the samples separate clearly per cell stage when taking the 5,000 most variable peaks into account (Figure 2C). Performing a two-group comparison between the pre-T and T stage samples, reveals 5,079 and 2,785 differentially accessible regions (false discovery rate, FDR < 1e-5, abs(log2FC) > 1) specific to the pre-T and T stage, respectively (Figure 2D). Aggregating the accessibility scores per sample over the differentially called regions visually corroborates the comparison (preT vs T) that was made (Supplemental figure S4). This is further illustrated by visualizing two called differential loci (Supplemental figure S5). In line with known aspects of thymocyte development (26), motif enrichment analysis reveals an expected enrichment pattern of the Runx1 transcription factor motif (cluster RUNX1,RUNX2,RUNX3) and the Mef2c transcription factor motif (cluster MEF2A,MEF2B,MEF2C) in peaks specific to the pre-T stage, while the LEF1 transcription factor motif (cluster LEF1,TCF7,TCF7L1) density is higher in the T stage (Supplemental figure S6). Complementary to the two-group comparison, an LRT analysis was performed for the cell type covariate. This analysis revealed clusters of peaks specific to the DN1, DN2/DN3, DN2/DN3/DP, DP, CD4 and CD4/CD8 stages (Figure 2E). Furthermore, patterns of motif enrichment across these clusters confirms the observations made in the two-group comparison, but reveal a more detailed pattern. For example, enrichment of the LEF1 transcription factor motif is mainly driven by peaks specific for CD4 cells and similarly Runx1 transcription factor motif enrichment can mainly be attributed to peaks from DN1 cells Supplemental figure S7). Both results highlight the use case for both the two-group mode as well as the LRT mode as both results are complementary though provide insights at different levels of granularity.

**Figure 2.**
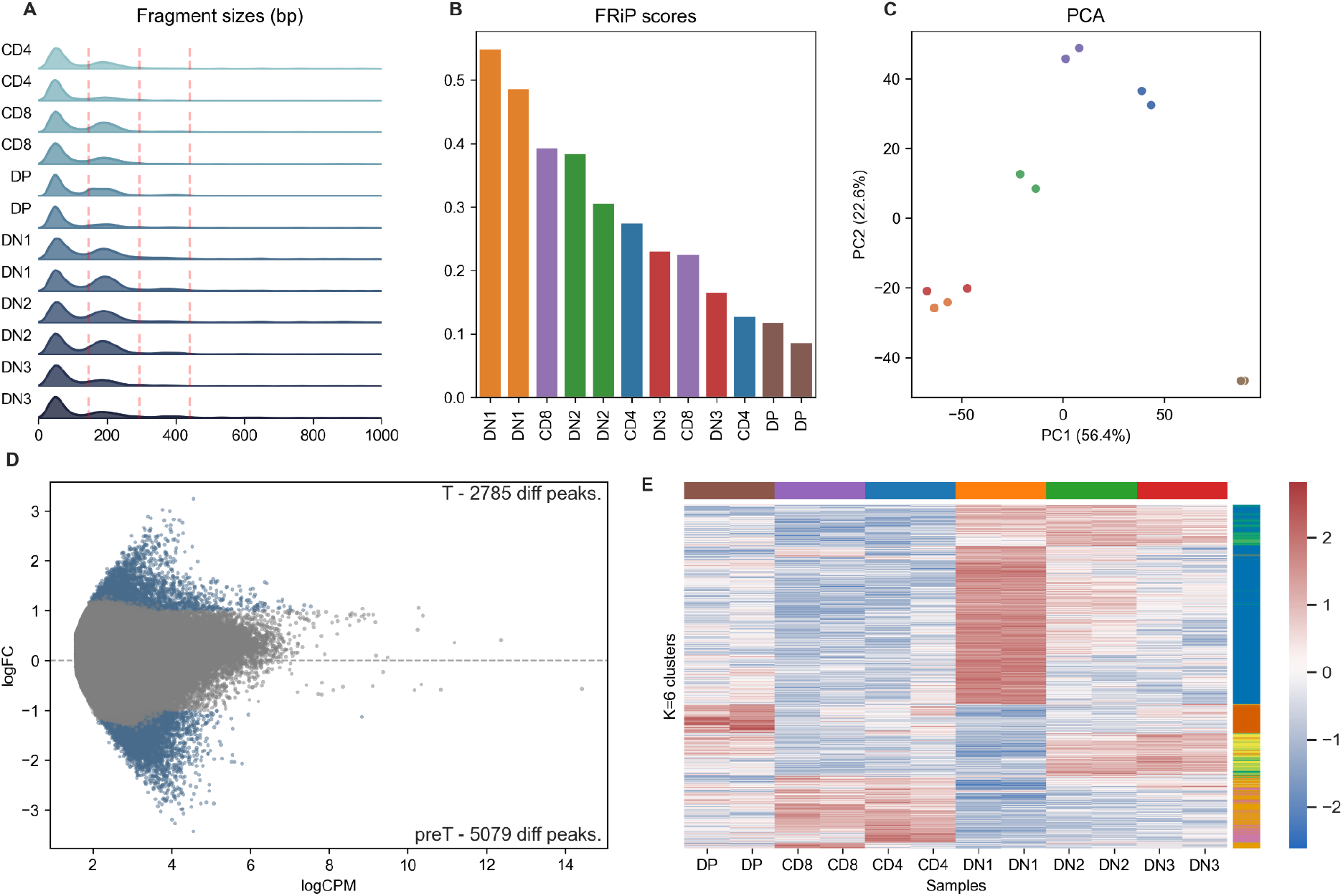
Overview of the generated quality metrics and differential accessibility scores for the ImmGen dataset. (A) Kernel density estimations for the fragment size distribution of all included samples. Nucleosomal bounds are indicated with dashed red lines at 147 bps (mononucleosomal), 294 bps (dinucleosomal) and 441 bps (trinucleosomal). (B) The fraction of reads in peaks (FRiP) scores calculated over the union of all peaks detected are shown per sample. (C) Sample grouping per cell type evaluated in a PCA plot. The percentage of variance explained per PC is indicated on the axis labels. (D) MAplot for the pre-T versus T two-group comparison. The number of differential peaks (FDR < 1e-5, abs(log2FC) > 1) per group are denoted on the plot and highlighted in blue. (E) Heatmap denoting the row-wise Z-scores of peaks significant (FDR < 1e-10) in the LRT analysis. Rows denote peaks and columns denote samples. Peaks clustered together by K-means clustering (K=6) are indicated by the vertical color bar. Sample color coding in the PCA plot (C) and the heatmap (E) follow the scheme set in (B).

### ATACofthesnake enables continuous time course analysis

Additional to the two-group and LRT modules, AOS allows for time course analyses as well. This is centered around Gaussian process regression (see methods), and AOS implements solutions where time is treated either continuously or as an ordinal covariate. As a proof-of-concept, time series ATAC-seq data derived from *C. elegans* by Gaidatzis *et al*. (27) was used. This data consists of developmental time points achieved from synchronized larvae, and thus allows the time covariate to be encoded continuously. Furthermore, there is both a treatment arm (auxin treated, to deplete *grh1* via a degron system) and a mock arm (EtOH treated). The standard QC module demonstrates high signal-to-noise ratios across the entire time course with FRiP scores ranging between 0.28 and 0.54, indicating that the data is of good quality (Figure 3A). The vast majority of between-group variability can be accounted for by the time-covariate, in contrast to the treatment arms (Figure 3B). After fitting a Gaussian process regression on all peaks, an empirical p-value was calculated by permuting the time covariate 1,000 times (see methods). In total, 8,930 peaks with a significant time signal (FDR < 0.01) could be identified. Subsequent k-means clustering of the time sensitive peaks reveals six groups with a distinct accessibility pattern over time (Figure 3C). The peaks underlying the clusters likely represent a conservative subset of all truly dynamic peaks, as can be observed through the clear aggregation of small uncorrected empirical p-values calculated for all peaks (Figure 3D). When visualizing normalized accessibility scores for a single peak per sample against predicted values of their corresponding fitted model, it is clear that the approach taken to identify time-dependent peaks is sensitive enough to model changes over small increments of time (Figure 3E-G), all while being robust enough to not overfit patterns on peaks that do not contain a clear temporal signal (Figure 3H). Furthermore, the ability to directly test the interaction between time and treatment allows subsequent identification of time-dependent peaks that display an altered temporal pattern in a specific treatment arm without assuming a specific directionality or structure (Supplemental figure 8). Taken together, the implementation of Gaussian process regression within AOS provides a framework to identify, characterize and group time-dependent peaks, across and between experimental conditions.

**Figure 3.**
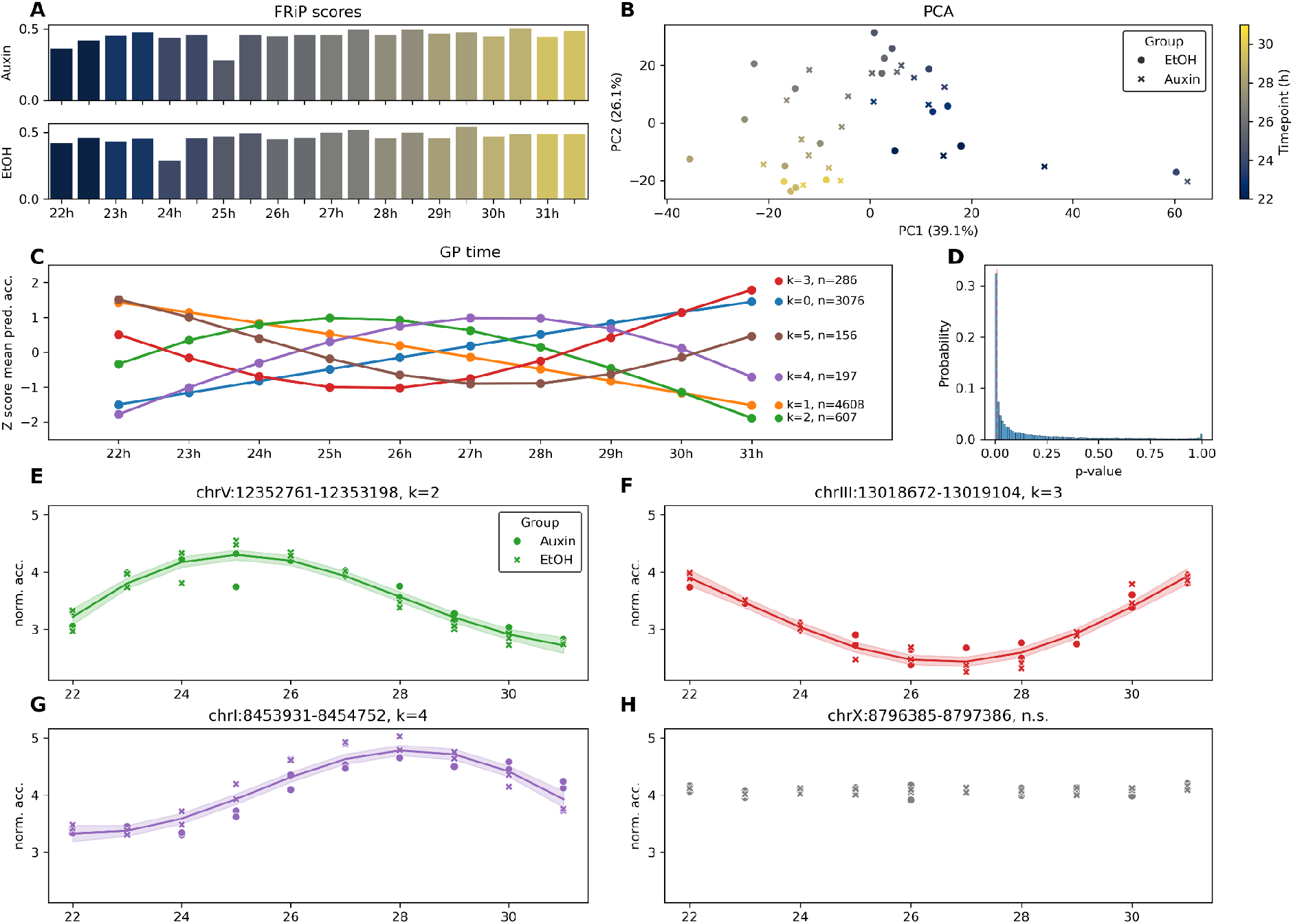
Overview of the generated quality metrics and time course results for the Gaidatzis dataset. (A) FRiP scores calculated over the union of all peaks detected per sample, either for the Auxin treated samples (top) or the mock (EtOH) treated samples (bottom). Color intensity and the x-axis denote the timepoint. (B) Grouping of the samples visualized in a PCA plot, where the symbol denotes the sample group (either Auxin treated or EtOH treated), and the color intensity denotes, similar to (A), the timepoint in hours. (C) Average trajectories (calculated as the mean of the row-wise Z scores per cluster, per timepoint) for the 5 clusters of time-dependent peaks. The number of peaks per cluster is indicated next to the respective curves. (D) The density of (uncorrected) p-values for the likelihood permutation test is indicated, with a vertical dashed red line indicating the FDR cutoff (1e-2). (E-H) Predicted values from the fitted data for a single member (peak) of that cluster (k=2 (E), k=3 (F), k=4 (G), time-independent peak (H)). The data points denote the normalized accessibility scores, the line denotes the posterior predictive mean while the shaded area denotes the posterior standard deviation. The circles and crosses indicate the normalized accessibility values from the original samples derived from the Auxin group, or the mock treated group, respectively.

### ATACofthesnake enables time course analysis in an ordinal setting

To exemplify the use of AOS in a setting where time is treated as an ordinal covariate, the ImmGen data was revisited. Given that this dataset comprises a differentiation trajectory of thymocytes, the samples could be seen as originating from ordinal time points. Doing so reveals, similar to the continuous time course analysis, six groups of peaks with a specific temporal pattern over the differentiation course (Figure 4A). Additionally, the distribution of uncorrected p-values hints (similar to what was seen in the continuous setting, Figure 3D) that the majority of accessible sites are dynamic over time (Figure 4B). Just as in the continuous case, individual fitted curves match the accessibility scores particularly well (Figure 4C,D), indicating that the Gaussian process regression approach is well suited for the ordinal case too. Additionally, rather than equidistantly spacing the ordinal time-points, the distances between the time points are treated as hyperparameters and are estimated during the fitting of the model too. By estimating these ‘pseudodistances’ for each peak, an indication of where along the ordinal time trajectory the largest change in accessibility occurred is obtained. Interestingly, clustering the data based on the pseudodistances instead of the inferred temporal patterns, reveals six clusters. Four of those have a single dominant transition point where the accompanying change in accessibility is largest between the two adjacent time points (Figure 4E). Furthermore, peaks assigned to the pseudodistance-based clusters were distributed across all previously identified temporal groups, indicating that clustering based on pseudodistances captures an additional aspect that would be difficult to obtain otherwise (Figure 4F). These patterns hold true for all clusters, and are also the case for the two additional clusters where the transition is dominant over two time points instead of just one (Supplemental figure S9). Taken together, these findings show that the estimated pseudodistances can reveal an additional layer on the temporal structure over the entire time course by identifying the intervals in which accessibility changes are most pronounced.

**Figure 4.**
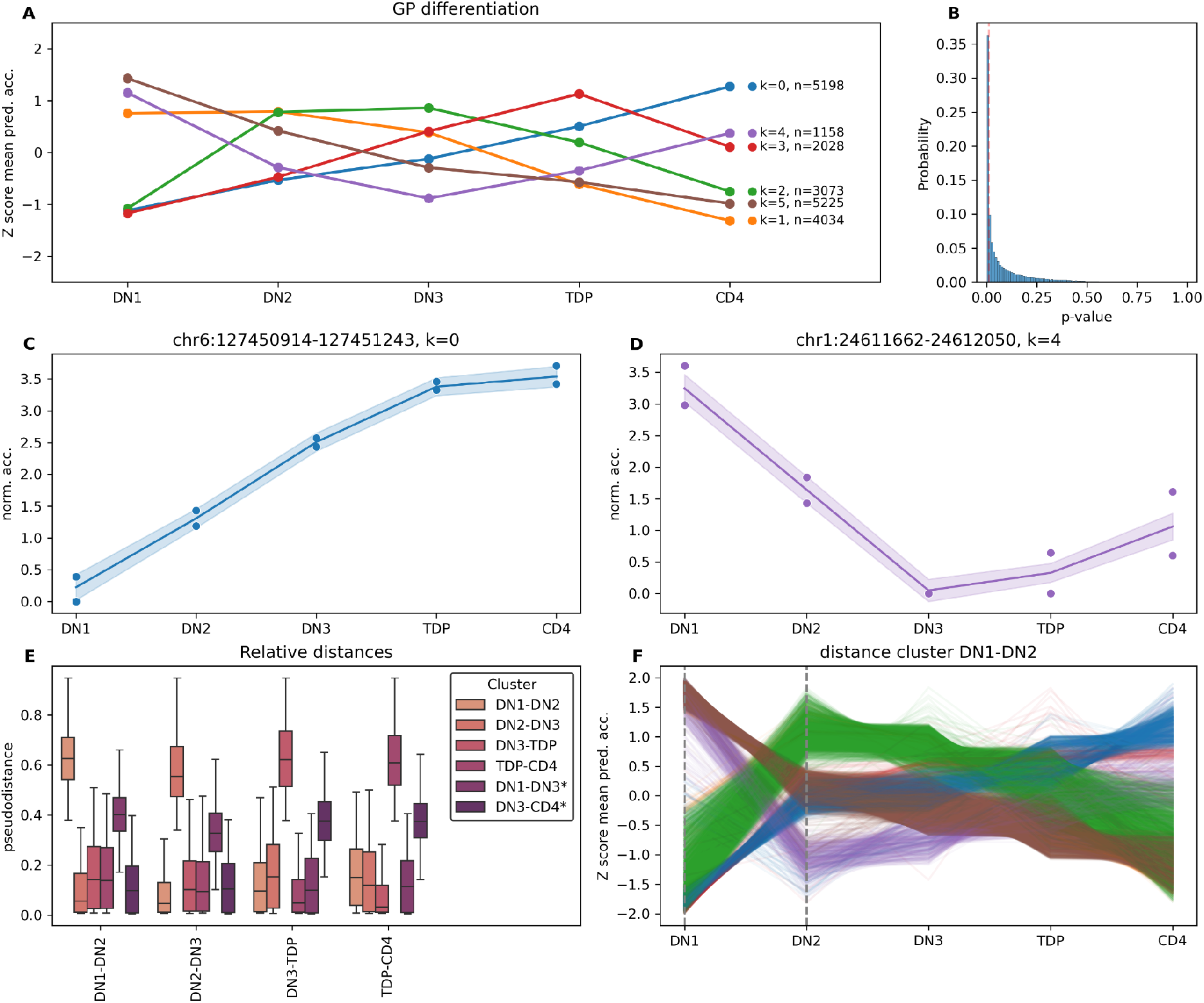
The ImmGen data treated as an ordinal time course experiment. (A) The average trajectories (calculated as the mean of the row-wise Z scores per cluster, per differentiation stage) for the 6 clusters of time-dependent peaks. The number of peaks per cluster is indicated next to each curve. (B) Density of (uncorrected) p-values for the likelihood permutation test. The vertical dashed red line indicates the FDR cutoff (1e-2). (C-D) Predicted values from a single member (peak) for cluster k=0 (C) and cluster k=4 (D) indicated as colored lines with the shaded area denoting the posterior standard deviation of the predicted mean. The normalized accessibility values per sample are indicated on the curves as dots. (E) For every stage transition, the distribution of estimated pseudo distances per peak are indicated, split up per identified cluster. (F) The individual trajectories for single peaks belonging to cluster DN1-DN2 are denoted as Z scores, with the color of the individual trajectories matching their cluster as determined by the temporal patterns seen before (A).

### ATACofthesnake automatically performs motif enrichment and footprinting analyses after differential mode is requested

If any of the aforementioned differential modes are requested (either two-group, LRT and/or time course analyses), automatic postprocessing (motif enrichment and footprinting analyses) on the significant results is performed when a motif file is provided. Motif enrichment is only performed if sufficient significant peaks are identified (the threshold for ‘sufficient’ is user-parametrizable) and subsequently footprinting analysis is only performed if significantly enriched motifs can be scored to begin with. The FDR cut-off to be used in motif enrichment can be customized by the end user as well. The added value of this analysis can be illustrated while revisiting the ordinal time course analysis on the ImmGen data, by zooming in on two enriched motifs (LEF1 and RUNX1), identified in cluster k=0, clusters k=1 and k=2, respectively (Figure 4A). Aggregating the accessibility scores over motif occurrences reveals that for both motifs, the accessibility patterns around the motifs follow the average accessibility pattern of the cluster itself. Furthermore, the actual motif sites reveal a relative drop in accessibility at specific time points, indicating that they are effectively bound at these stages (Figure 5). Taken together, AOS comprises a framework that enables differential accessibility analysis by allowing both simple, complex and time-course analyses all while facilitating drawing biological conclusions by automating motif enrichment and footprinting analyses.

**Figure 5.**
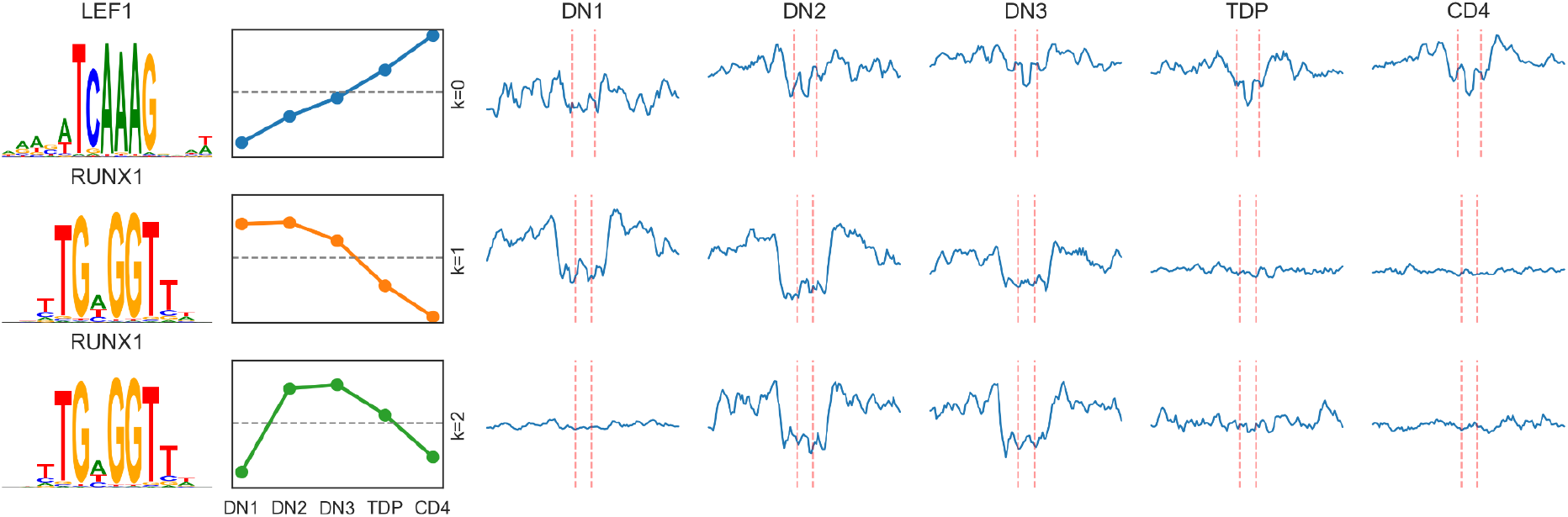
Footprinting analyses results for two enriched motifs found in the ImmGen data treated as an ordinal time course experiment. Indicated are two found enriched consensus motifs (left), the average accessibility scores for a specific cluster (middle), and the corrected accessibility scores per ordinal timepoint centered around motif occurrences (denoted in dashed red lines, right).

## Discussion

In this work, a versatile framework is presented to analyze multifactorial chromatin accessibility data. Given its emphasis on the analysis of experiments with a complex design, AOS provides a scalable solution that handles large datasets with ease. Furthermore, since all covariates denoted in the samplesheet are used in the subsequent models, it becomes trivial to combine data from different studies or sequencing sources, as these can be encoded as nuisance variables. Ease of use was a central theme in the planning and design of the workflow, and the fact that requesting a specific comparison is done on a sample level rather than on a coefficient level makes encoding complex designs straightforward. To ensure the framework adheres to FAIR principles as much as possible, AOS automatically logs all configuration parameters, runtime settings, and data paths, guaranteeing that analyses are fully transparent. The availability of both two-group comparisons and LRT analyses makes that for some analyses, complementary solutions are available but could also aid the discovery of new biological insights that would otherwise be missed. The addition of the time course mode adds to this complementarity, as for example time course analyses could also be performed using an LRT analysis. The advantage of having multiple options is that they each have specific statistical constraints. Using Gaussian process regression, for example, allows to capture continuity better than encoding time as a categorical covariate in an LRT analysis. Additionally, the option to perform ordinal time course analyses is a feature that to date is not easily performed on chromatin accessibility data. By estimating pseudodistances between the ordinal time points, a new way to classify peaks emerged. By revisiting the ImmGen dataset, both the two-group mode and LRT analyses could be validated, as expected motifs known to be important in murine thymopoiesis could be retrieved. Revisiting the Gaidatzis *et al*. dataset allowed us to test the time course mode in a continuous fashion, and doing so revealed five clusters with specific time trajectories that could be modeled in an accurate manner. The corroboration of biological insights and consistent results on real-world data is promising, though a number of connotations can still be made. First of all, AOS is built on snakemake, which enables parallel processing of data, as utilizing high-performance compute instances becomes effortless. Complex experimental design and the number of samples are directly correlated, and AOS can handle many samples effortlessly due to the snakemake backend. Nevertheless, since determination of significance in the time course mode is based on permutation, runtimes for experiments with many peaks need to be taken into consideration. Even though these are still measured in hours rather than days, reducing the number of permutation iterations could alleviate this manner, or one could switch to an LRT analysis. All-in-all, AOS comprises a versatile framework, with multiple options to determine differential accessible regions and downstream analysis in a general way, making it an optimal go-to for most chromatin accessibility datasets.

## Conclusion

With AOS, a gap in the analysis of chromatin accessibility data is bridged by providing a comprehensive, statistically rigorous framework that handles multifactorial data seamlessly. The modular design and implementation in snakemake makes the analysis FAIR and intuitive for both experimentalists and bioinformaticians alike. Implementation of two-group and LRT analyses tackles most common use cases, and the realization of Gaussian process regression for both continuous and ordinal time course analyses brings a novel technique to typical analyses done on chromatin accessibility data. The validation on real-world datasets demonstrates the workflow’s capacity to recapitulate known biological insights while enabling discovery of patterns that would be difficult to detect otherwise. As chromatin accessibility profiling continues to grow in both complexity and scale, AOS provides the community with a scalable, flexible, and statistically sound solution for extracting biological meaning from multifactorial experimental designs.

## Methods

### Implementation

AOS is a modular workflow written in snakemake (19). In the default mode, aligned reads overlapping with read-attracting regions (defined by user-input) are removed and only reads from putative nucleosome free regions (with fragment sizes below 150 bps) are retained using alignmentSieve implemented in deepTools (28). From the original BAM and/or CRAM files, fragment size distributions and reads overlapping with the mitochondrial genome are determined using bamPEFragmentSize implemented in deepTools and samtools (29), respectively. Sieved BAM files are subsequently converted into bed files using bedtools (30) and used to call peaks using MACS3 (31), using ‘-f BED --nomodel --shift -75 --extsize 150 - q 0.01’ as flags. This effectively takes all reads into consideration, and ensures that the peak signal is centered around Tn5 cutsites. Next, the union of the called peaks across all samples is determined with bedtools merge, and a count matrix is generated with multiBamSummary implemented in deepTools. Peak to feature annotation is performed on the unionized peakset with UROPA (32), and fraction of reads in peaks per sample metrics are calculated with samtools. Bigwig files (both RPKM and scalefactor normalized) are made for visualization using the bamCoverage function implemented in deepTools. Quality metric visualisation and the visualisation of results in default mode (and differential mode) are performed with matplotlib (33) and seaborn (34). The differential accessibility mode consists of three modules, two of which (the two-group module and LRT module) use edgeR at their core (24). Pairwise differential accessible peaks between two conditions are determined in the two-group module using the quasi-likelihood framework available in edgeR (glmQLFTest function), while peaks significant under an LRT test are determined using the glmLRT function available in edgeR. The end-user is responsible for formulating a reduced design. If a sufficient number (defined by the end-user) of peaks are significant under the LRT test, K-means clustering is performed on Z-score transformed counts of those peaks. The optimal number of clusters (K) is determined automatically by maximizing the distance between the line perpendicular to the inertia values and the line connecting the first (K=2) and last (K=19) inertia values. K-means clustering is performed using scikit-learn (35).

### Time course analysis

Raw count matrices are first CPM (counts per million) normalized and log-transformed:

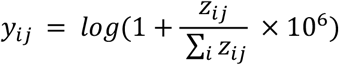

With *z* denoting the raw counts for peak *i* in sample *j*. Subsequently, for each peak independently, a Gaussian process regression model is fitted to model chromatin accessibility over time:

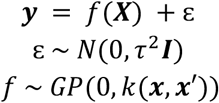

with τ^2^ = 0.1 by default, but customizable via the command line.The actual kernel *k* depends on what time course analysis is requested. If time is treated as a continuous covariate, the kernel *k* is set as a radial basis function kernel (RBF) with automatic relevance determination (ARD) (via the *l*_*d*_- hyperparameter):

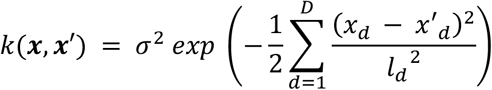

with both *σ* and *l* being optimized from the data using five random, independent restarts. *D* denotes the input covariates, including time. An additional model is fitted, where the time covariate is omitted (***X***_*D-*1_) and the log marginal likelihood ratio is calculated:

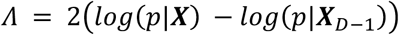

Significance testing is done by randomly permuting the time covariate *N* times, and subsequently calculating an empirical p-value with:

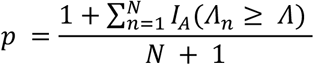

with *I*_*A*_ denoting an indicator function and *Λ*_*n*_ the log marginal likelihood ratio for the permuted models. The p-values are subsequently corrected with the Benjamini-Hochberg procedure.

Covariate-time interactions can be tested as well, which follows a similar Gaussian process regression scheme, with the difference that the covariate kernel is decomposed into a time kernel (*k*_*t*_), a covariate of interest kernel (*k*_*z*_) and (when relevant) a kernel with nuisance covariates (*k*_*v*_), with ***X*** = (***t, z*, *v***). The kernel function is subsequently defined as:

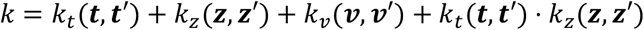

Obtaining the log marginal likelihood ratio and the subsequent empirical p-value is done in a similar fashion, by removing the product kernel encoding the interaction term.

In the ordinal setting, the starting point is again the CPM normalized, log-transformed count matrix. Additionally, the unique time points (*T*) are treated as ordered categories where subsequent time points are strictly larger than the previous one:

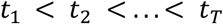

To determine the pseudodistances between time points, a set *α* of *T* − 1 parameters are defined to represent the spacings *s* between the time points, using a softmax transformation:

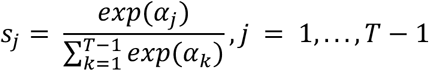

such that the sum of all spacings is 1. Next, a warped time variable can be defined as:

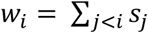

Initially, uniform spacing is assumed between the time points. Subsequently, the spacing parameters are learned from the data by maximizing the marginal likelihood jointly with the other kernel hyperparameters. The entire implementation is based on the GaussianProcessRegressor functionality available within scikit-learn (35).

### Postprocessing of significant peaks

Regardless of the module used to generate differential peaks (two-group -, LRT - or time course mode), postprocessing of significant peaks (which can be parametrized both with a number cutoff and an FDR cutoff) can be done if a list of motifs is provided. To this extent, the ame functionality implemented in the MEME suite (36) is used to score enriched motifs for every comparison calculated. This is done on a per-group basis. More specifically, motif enrichment is performed for every group (either one of the two groups found in the two-group module, or decided by K-means clustering in case of the LRT or time course module) using the same set shuffled as background. Additionally, all replicates within a group are subsampled to the same depth and merged together to perform footprinting analysis using TOBIAS (17). Both ATAC corrected and footprinting scores are generated using the ATACorrect and ScoreBigwig functions, and for every significant motif found in the motif enrichment, individual instances are scored using fimo tool (available in the MEME suite) and the aggregated accessibility signal visualized using the plotAggregate function (available in TOBIAS).

### Public dataset analysis

Both the ImmGen dataset (25) and the Gaidatzis dataset (27) were processed using snakePipes (version 3.2.0) using the --dedup, --fastqc and --trim flags. References used for alignment were the GRCm39 genome reference with ENSEMBL version 106 for the ImmGen dataset, and the wormbase PRJNA13758 reference with WS285 based gene models for the Gaidatzis dataset. The resulting deduplicated bam files were directly used as input for ATACofthesnake (version 0.13.0). All data used to generate figures and metrics were generated by AOS, with the exception of the single locus accessibility scores (Supplemental figure S5), which was generated using pyGenomeTracks (37).

## Data availability

The datasets analysed in this study are publicly available under GEO accessions GSE100738 (ImmGen data) (25) and GSE288914 (Gaidatzis data) (27). The processed data of these accessions (used as input for the workflow), as well as all metadata such as samplesheets, comparison files, gene models and reference genome sequences is made available via zenodo under accession 19296250 (38). Downloading the example data and metadata is automated and can be performed by using the ‘example’ functionality provided within the AOS workflow. ATACofthesnake is publicly available via github (https://github.com/maxplanck-ie/ATACofthesnake) and can be installed using PyPi (https://pypi.org/project/ATACofthesnake/). Usage documentation is available under ReadTheDocs (https://atacofthesnake.readthedocs.io/). The code used to generate the figures in this manuscript is documented and available via github (https://github.com/WardDeb/aos_example).

## Author contributions

WD conceptualized the study, wrote the software and the manuscript, and analyzed the data. AL conceptualized the study, interpreted the biological results, wrote the manuscript and analyzed the data. AS wrote the software. TM conceptualized and supervised the study. All authors read and approved the manuscript.

## Acknowledgements

We would like to thank early adopters of the AOS workflow, for valuable feedback and for providing user experiences. In particular Nikolay Zolotarev, Arion Foertsch, Sarah Bowden and Galina Erikson. We acknowledge project support from the Max Planck Computing and Data Facility. We acknowledge computing support from the Scientific Data Processing Unit from the Max Planck Institute of Immunobiology and Epigenetics, in particular support from Wolfgang Burger and Christian Pagel.

## References

1. Klemm SL, Shipony Z, Greenleaf WJ. Chromatin accessibility and the regulatory epigenome. Nat Rev Genet. 2019 Apr;20(4):207–20.

2. Hamrud E, Leese J, Thiery AP, Buzzi AL, Vigilante A, Briscoe J, et al. Dynamic changes in chromatin accessibility during cell fate specification at the neural plate border. Development [Internet]. 2025 Dec 1;152(23). Available from: http://dx.doi.org/10.1242/dev.204943

3. Trevino AE, Sinnott-Armstrong N, Andersen J, Yoon SJ, Huber N, Pritchard JK, et al. Chromatin accessibility dynamics in a model of human forebrain development. Science. 2020 Jan 24;367(6476):eaay1645.

4. Rispal J, Escaffit F, Trouche D. Chromatin dynamics in intestinal epithelial homeostasis: A paradigm of cell fate determination versus cell plasticity. Stem Cell Rev Rep. 2020 Dec;16(6):1062–80.

5. Scott-Browne JP, López-Moyado IF, Trifari S, Wong V, Chavez L, Rao A, et al. Dynamic changes in chromatin accessibility occur in CD8+ T cells responding to viral infection. Immunity. 2016 Dec 20;45(6):1327–40.

6. Waag R, von Ziegler L, Sonder E, Sturman O, Leonardi J, Frei S, et al. Distinct mechanisms of transcriptomic habituation to repeated stress in the mouse hippocampus. Nat Commun. 2025 Dec 22;16(1):11569.

7. Li Z, Zhang M, Zhang Y, Gan Y, Zhu Z, Wang J, et al. Integrative analysis of gene expression and chromatin dynamics multi-omics data in mouse models of bleomycin-induced lung fibrosis. Epigenetics Chromatin. 2025 Mar 12;18(1):11.

8. Bergo V, Bousounis P, To Vu G, Douté M, Polyzou A, Lalioti ME, et al. Lack of MDA5 delays hematopoietic aging by modulating inflammaging and proteostasis in mice. Nat Commun. 2026 Feb 12;17(1):1645.

9. Minnoye L, Marinov GK, Krausgruber T, Pan L, Marand AP, Secchia S, et al. Chromatin accessibility profiling methods. Nat Rev Methods Primers. 2021 Jan 21;1(1):10.

10. Buenrostro JD, Wu B, Chang HY, Greenleaf WJ. ATAC-seq: A method for assaying chromatin accessibility genome-wide. Curr Protoc Mol Biol. 2015 Jan 5;109(1):21.29.1– 21.29.9.

11. Corces MR, Trevino AE, Hamilton EG, Greenside PG, Sinnott-Armstrong NA, Vesuna S, et al. An improved ATAC-seq protocol reduces background and enables interrogation of frozen tissues. Nat Methods. 2017 Oct;14(10):959–62.

12. Schmid K, Schenk RP, Wiedemann GM. Low-input assay for transposase-accessible chromatin identifies epigenetic signatures of liver group 1 innate lymphoid cells. Eur J Immunol. 2025 Oct;55(10):e70066.

13. Hohl T, Bönisch U, Manke T, Arrigoni L. Enhancing single-cell ATAC sequencing with formaldehyde fixation, cryopreservation, and multiplexing for flexible analysis. BMC Res Notes. 2025 Oct 20;18(1):437.

14. Grandi FC, Modi H, Kampman L, Corces MR. Chromatin accessibility profiling by ATAC-seq. Nat Protoc. 2022 June;17(6):1518–52.

15. Barnett KR, Decato BE, Scott TJ, Hansen TJ, Chen B, Attalla J, et al. ATAC-Me captures prolonged DNA methylation of dynamic chromatin accessibility loci during cell fate transitions. Mol Cell. 2020 Mar 19;77(6):1350–64.e6.

16. Lenaerts A, Kucinski I, Deboutte W, Derecka M, Cauchy P, Manke T, et al. EBF1 primes B-lymphoid enhancers and limits the myeloid bias in murine multipotent progenitors. J Exp Med. 2022 Nov 7;219(11):e20212437.

17. Bentsen M, Goymann P, Schultheis H, Klee K, Petrova A, Wiegandt R, et al. ATAC-seq footprinting unravels kinetics of transcription factor binding during zygotic genome activation. Nat Commun. 2020 Aug 26;11(1):4267.

18. Schep AN, Buenrostro JD, Denny SK, Schwartz K, Sherlock G, Greenleaf WJ. Structured nucleosome fingerprints enable high-resolution mapping of chromatin architecture within regulatory regions. Genome Res. 2015 Nov;25(11):1757–70.

19. Mölder F, Jablonski KP, Letcher B, Hall MB, van Dyken PC, Tomkins-Tinch CH, et al. Sustainable data analysis with Snakemake. F1000Res. 2021 Jan 18;10:33.

20. Wilkinson MD, Dumontier M, Aalbersberg IJJ, Appleton G, Axton M, Baak A, et al. The FAIR Guiding Principles for scientific data management and stewardship. Sci Data. 2016 Mar 15;3(1):160018.

21. Bhardwaj V, Heyne S, Sikora K, Rabbani L, Rauer M, Kilpert F, et al. snakePipes: facilitating flexible, scalable and integrative epigenomic analysis. Bioinformatics. 2019 Nov 1;35(22):4757–9.

22. Ewels PA, Peltzer A, Fillinger S, Patel H, Alneberg J, Wilm A, et al. The nf-core framework for community-curated bioinformatics pipelines. Nat Biotechnol. 2020 Mar;38(3):276–8.

23. Ewels P, Käller M. MultiQC: visualising results from common bioinformatics tools [Internet]. 2017. Available from: http://dx.doi.org/10.7490/f1000research.1114455.1

24. Robinson MD, McCarthy DJ, Smyth GK. edgeR: a Bioconductor package for differential expression analysis of digital gene expression data. Bioinformatics. 2010 Jan 1;26(1):139–40.

25. Yoshida H, Lareau CA, Ramirez RN, Rose SA, Maier B, Wroblewska A, et al. The cis-regulatory atlas of the mouse immune system. Cell. 2019 Feb 7;176(4):897–912.e20.

26. Hosokawa H, Rothenberg EV. How transcription factors drive choice of the T cell fate. Nat Rev Immunol. 2021 Mar;21(3):162–76.

27. Gaidatzis D, Graf-Landua M, Methot SP, Wölk M, Brancati G, Hauser YP, et al. A scheduler for rhythmic gene expression. Mol Syst Biol. 2025 Dec;21(12):1793–821.

28. Ramírez F, Ryan DP, Grüning B, Bhardwaj V, Kilpert F, Richter AS, et al. deepTools2: a next generation web server for deep-sequencing data analysis. Nucleic Acids Res. 2016 July 8;44(W1):W160–5.

29. Li H, Handsaker B, Wysoker A, Fennell T, Ruan J, Homer N, et al. The Sequence Alignment/Map format and SAMtools. Bioinformatics. 2009 Aug 15;25(16):2078–9.

30. Quinlan AR, Hall IM. BEDTools: a flexible suite of utilities for comparing genomic features. Bioinformatics. 2010 Mar 15;26(6):841–2.

31. Feng J, Liu T, Zhang Y. Using MACS to identify peaks from ChIP-Seq data. Curr Protoc Bioinformatics. 2011 June;Chapter 2(1):2.14.1–2.14.14.

32. Kondili M, Fust A, Preussner J, Kuenne C, Braun T, Looso M. UROPA: a tool for Universal RObust Peak Annotation. Sci Rep. 2017 June 1;7(1):2593.

33. Hunter JD. Matplotlib: A 2D Graphics Environment. Comput Sci Eng. 2007 May;9(3):90– 5.

34. Waskom M. seaborn: statistical data visualization. J Open Source Softw. 2021 Apr 6;6(60):3021.

35. Buitinck L, Louppe G, Blondel M, Pedregosa F, Mueller A, Grisel O, et al. API design for machine learning software: experiences from the scikit-learn project [Internet]. arXiv [cs.LG]. 2013. Available from: 10.48550/arXiv.1309.0238

36. McLeay RC, Bailey TL. Motif Enrichment Analysis: a unified framework and an evaluation on ChIP data. BMC Bioinformatics. 2010 Apr 1;11(1):165.

37. Lopez-Delisle L, Rabbani L, Wolff J, Bhardwaj V, Backofen R, Grüning B, et al. pyGenomeTracks: reproducible plots for multivariate genomic datasets. Bioinformatics. 2021 Apr 20;37(3):422–3.

38. Deboutte W. AOS example public dataset [Internet]. Zenodo; 2026. Available from: http://dx.doi.org/10.5281/ZENODO.19296250

