## Supplemental Tables and Figures for "ATACofthesnake: A scalable framework for analyzing multifactorial and time course chromatin accessibility data"

### Supplemental information

Supplemental table S1. Overview of required and optional arguments to run AOS.

| Argument | required | meaning |
|---|---|---|
| -i / --bamdir | **yes** | input directory containing input BAM / CRAM files |
| -o / --outputdir | **yes** | directory to write output to |
| -g / --gtf | **yes** | file containing the gene models |
| -r / --genomefasta | **yes** | reference genome |
| -b / --readattractingregions | **yes** | BED file containing mitochondrial genome and read attracting regions to omit |
| -p / --snakemakeprofile | no | snakemake profile to use schedulers / HPC submission |
| -@ / --threads | no | number of threads to use (if -p not set). Defaults to 1 |
| -m / --motifs | no | motif file in MEME format to run enrichment and footprinting analyses against |
| -f / --fragsize | no | Maximum fragment size to be considered in peak calling. Defaults to 150 bp. |
| --samplesheet | no | samplesheet (in tsv format) |
| --comparison | no | comparison (in yaml format) file to specific what differential analyses to be ran. |
| --mitostring | no | name of the mitochondrial genome contig. Defaults to ‘MT’ |
| --upstreamuro | no | maximum permitted distance upstream of a feature to be annotated to a peak. Defaults to 50000 |
| --downstreamuro | no | maximum permitted distance downstream of a feature to be annotated to a peak. Defaults to 50000 |
| --featureuro | no | feature to be considered in peak annotation (defaults to ‘gene’) |
| --pseudocount | no | pseudocount added to the count matrix prior to differential peak calling (two-group mode). Defaults to 8 |
| --peakset | no | external peak set (BED format) to be considered for count matrix generation |

| | | |
|---|---|---|
| --permutation_cutoff | no | p-value cutoff to determine significance testing after permutation (in time course mode). Defaults to 0.01 |
| --permutation_iterations | no | number of permutations to generate the p-value in time course mode. Defaults to 1000 |
| --fdr_cutoff | no | FDR cut-off to determine significance in two-group, LRT modes, and motif enrichment. Defaults to 0.001 |
| --lfc_cutoff | no | log2FoldChange cutoff used to determine significance in two-group mode (together with fdr_cutoff). Defaults to 0 |
| --min_sigpeaks | no | number of significant peaks required for a comparison to be considered for downstream analysis (motif enrichment and footprinting). Defaults to 100 |
| --gp_timesteps | no | number of steps in time course analysis for accessibility prediction. Defaults to 10 |
| --gp_alpha | no | the variance for the additional noise term in the time-course mode. Higher values result in more smoothing (flatter curves). Lower values result in less smoothing (more jagged fits). Defaults to 0.1 |

Supplemental figure S2. Benchmark results for the different processes executed. The top row represents a run on a dataset containing samples from T cell maturation process (ImmGen dataset), the bottom row represents a run with samples taken from a C. elegans time course dataset (Gaidatzis dataset). Rules from the default mode are indicated in orange, those from the differential mode are indicated in green. Rules that were executed multiple times (due to them being run on multiple input files) have their corresponding standard deviation indicated with the black errorbars. Both datasets were run with default settings (which includes 1000 permutations for the gaussian process regressions).

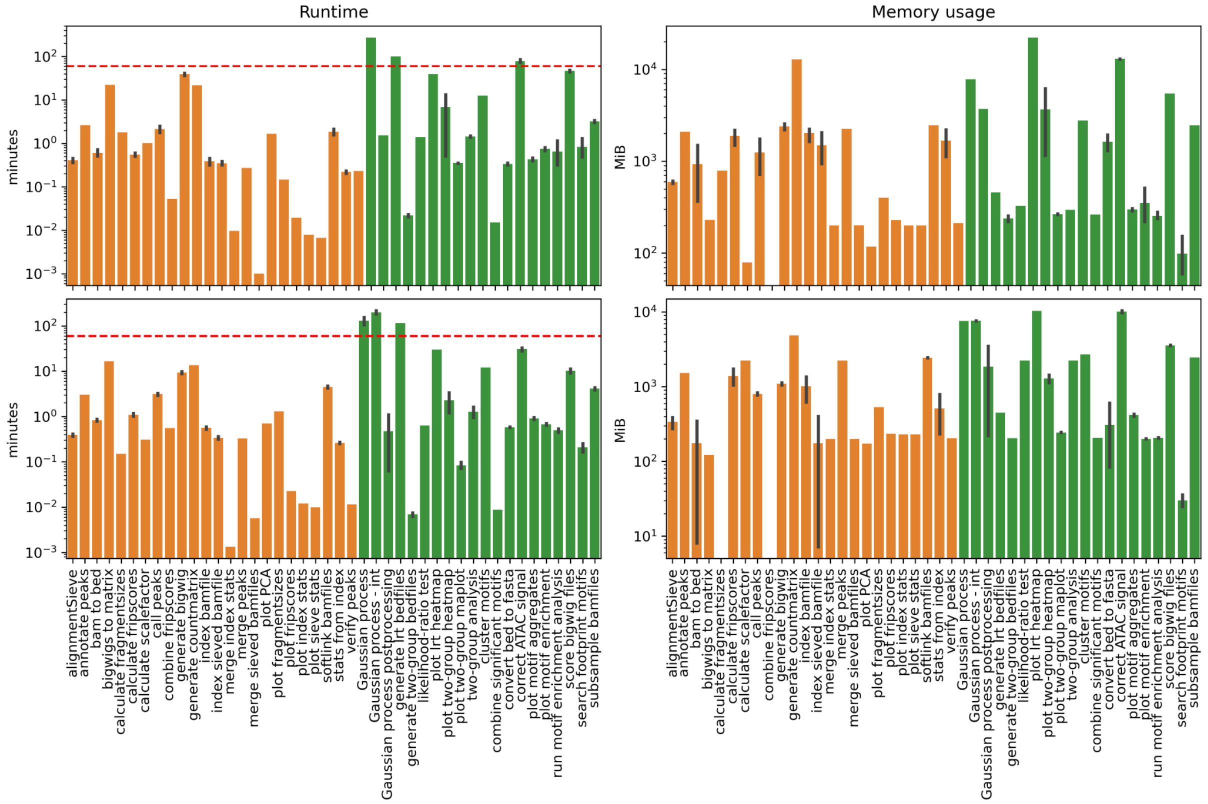

Supplemental table S3. Example of a samplesheet table and comparison yaml file for the ImmGen dataset. Two differential analyses are requested here in a single file, the first analysis requested is named ‘preT_vs_T’ is a two-group analysis comparing the pre-T stage (DN1, DN2, DN3) and the T stage (DP T cells, CD4 T cells and CD8 T cells), the second analysis is named ‘lrt_celltype’ and requests an LRT with an intercept only model as a reduced model.

Samplesheet:

| **sample** | **celltype** |
|---|---|
| DN1_rep1 | DN1 |
| DN1_rep2 | DN1 |
| DN2_rep1 | DN2 |
| DN2_rep2 | DN2 |
| DN3_rep1 | DN3 |
| DN3_rep2 | DN3 |
| CD4_rep1 | CD4 |
| CD4_rep2 | CD4 |
| CD8_rep1 | CD8 |
| CD8_rep2 | CD8 |
| DP_rep1 | DP |
| DP_rep2 | DP |

comparison.yaml:

```
preT_vs_T:
    type: ‘twogroup’
    preT:
      celltype:
          -   ‘DN1’
          -   ‘DN2’
          -   ‘DN3’
    T:
      celltype:
          -   ‘CD4’
          -   ‘CD8’
          -   ‘DP’
lrt_celltype:
    type: ‘lrt’
    reduced: ‘~1’
```

Supplemental figure S4. Differential accessible peaks called in the two-group comparison of the ImmGen dataset (preT vs T conditions). Columns denote different samples, rows denote the average aggregated signal per condition for T-specific peaks and pre-T specific peaks, respectively. Signal of the heatmap denotes normalized accessibility scores.

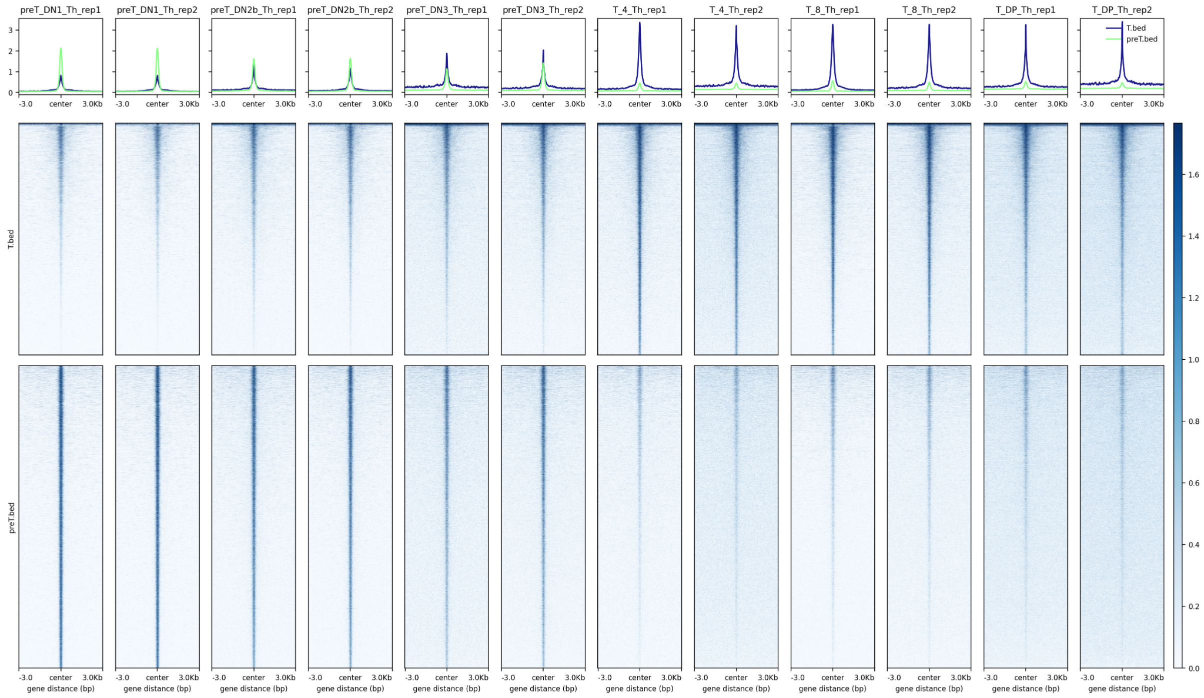

Supplemental figure S5. Examples of accessibility scores over two loci called in the preT vs T two-group comparison performed on the ImmGen dataset. The signal depicts scale factor normalized ATAC signal per sample. The top plot shows a peak specific to the preT stage (DN1, DN2 and DN3 cell types) over region chr1:52,157,360-52,201,703, while the bottom plot shows a peak specific to the T stage (CD4, CD8, DP) over region chr13:93,176,830-93,290,000. Peak boundaries for the differential peak are denoted by gray vertical lines.

### preT specific peak

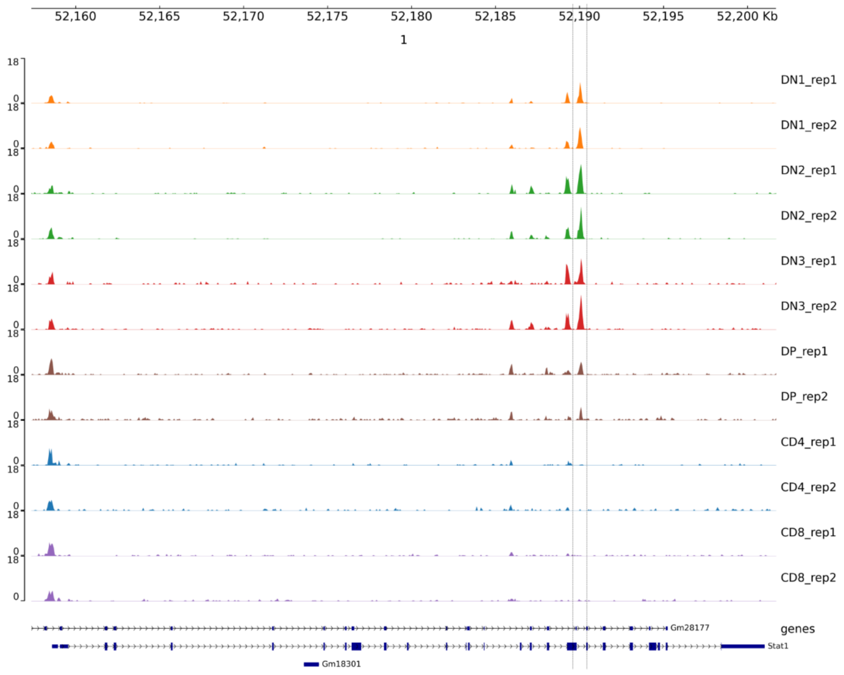


### T specific peak

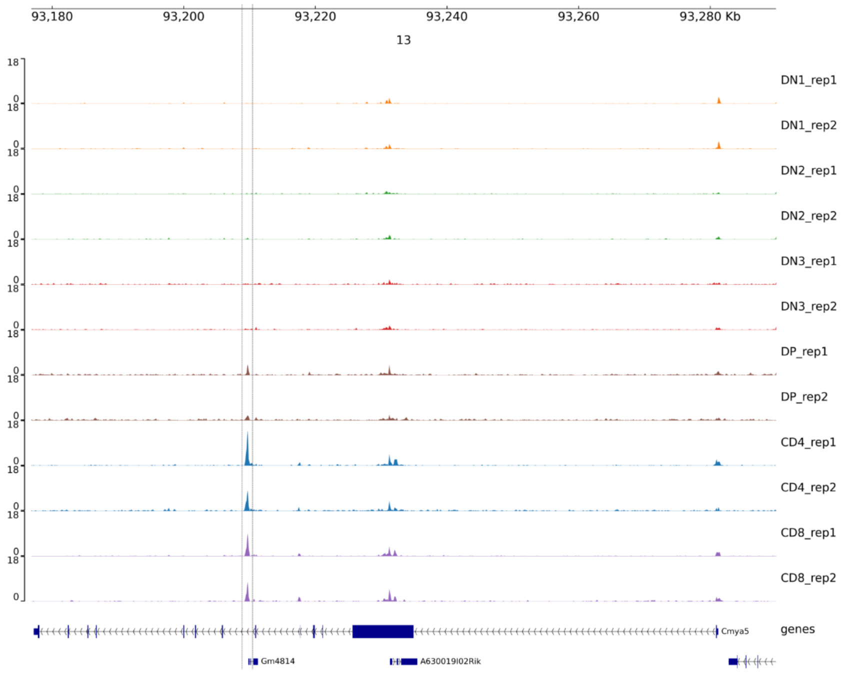

Supplemental figure S6. Motif enrichment results for the two-group comparison of the ImmGen dataset (preT vs T conditions). Only motifs that are significantly enriched (adjusted p-value < 1e-3) in at least one of the conditions were retained. The heatmap shows the motif density per kb, the single column heatmap shows the log2-transformed ratio of the densities per condition.

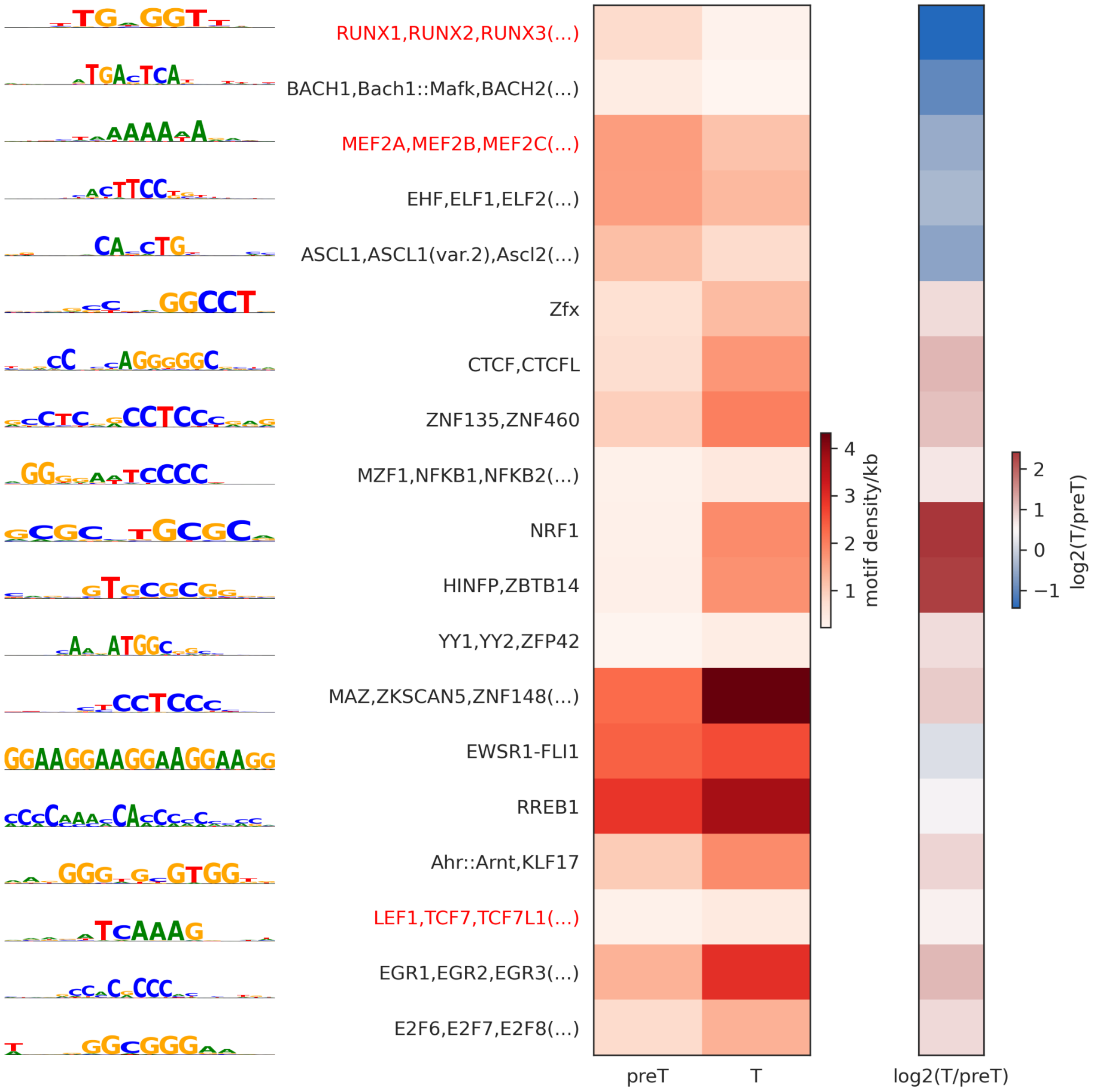

Supplemental figure S7. Motif enrichment results for the LRT analysis performed on the ImmGen dataset. The heatmap on the left shows Z scores of average accessibility values per K-means cluster (k=6). The second heatmap shows Z scores of motif densities per kb per cluster. Only motifs that are significantly enriched (adjusted p-value < 1e-3) in at least one of the conditions were retained.

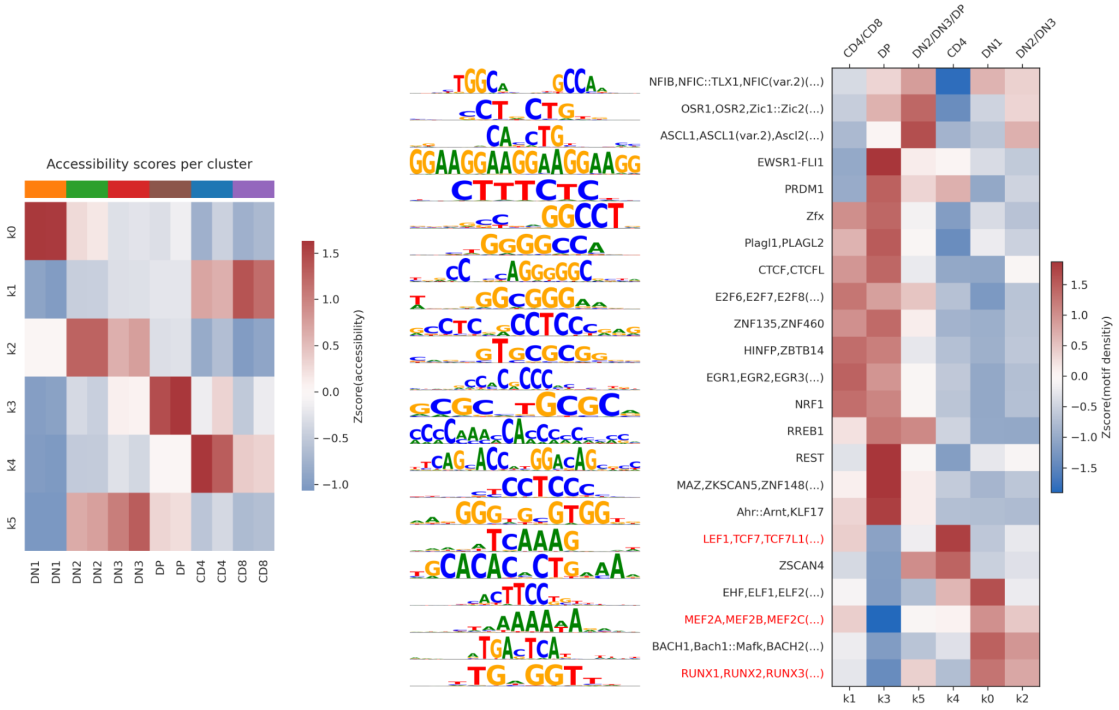

Supplemental figure S8. Example of three peaks showing an interaction effect for the Gaidatzis dataset. The three rows depict temporal patterns for three different genomic loci (peaks). Normalized accessibility scores for the Auxin arm are indicated in blue dots, while those from the EtOH treated samples are indicated with orange crosses. The blue and orange lines and shaded areas denote the fitted posterior means and posterior standard deviations for the Auxin arm and the EtOH arm, respectively.

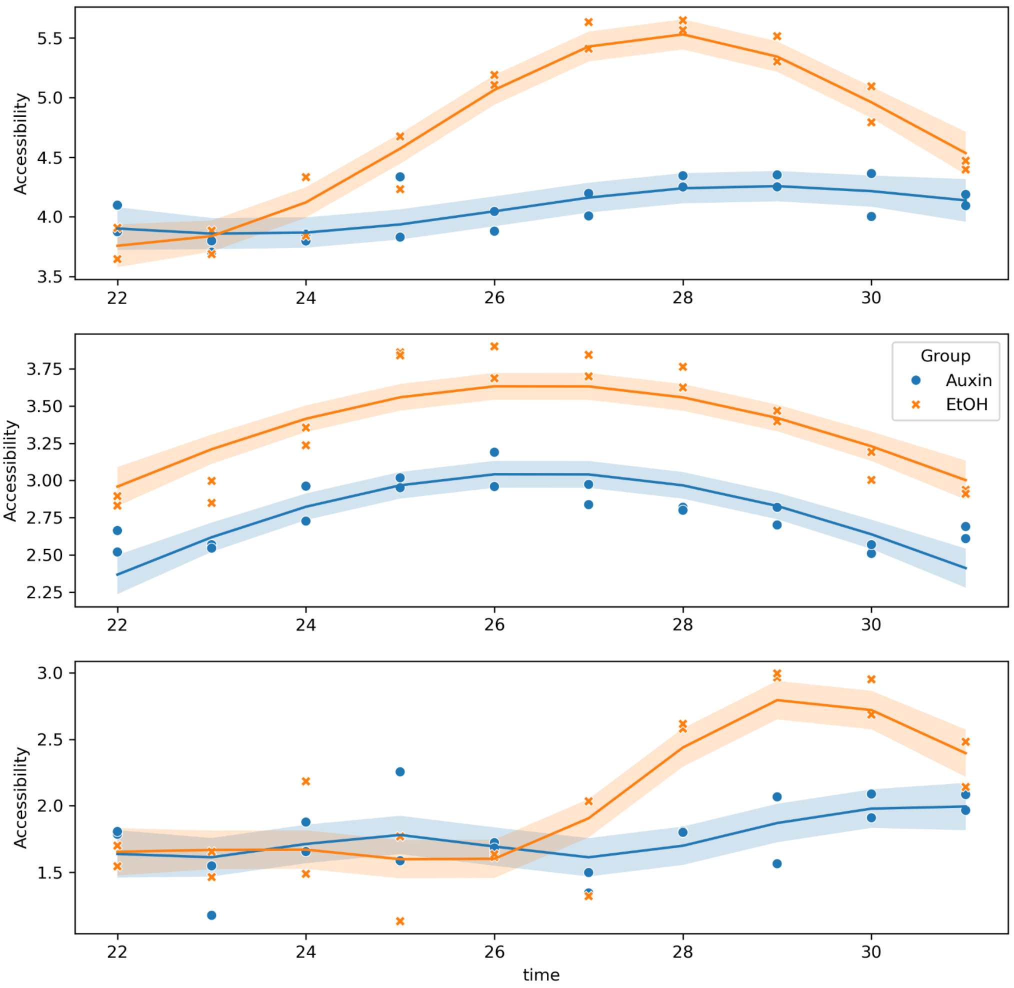

Supplemental figure S9. Time sensitive peaks identified using ordinal time course analysis are clustered based on their estimated pseudodistance. Every row denotes a cluster. The first two column depict the estimated pseudodistances individually (left) denoted together as lines where one line depicts a peak (middle). The temporal trajectories of the peaks underlying the pseudodistance-based clustering pattern are indicated as Z scores, where the color denotes the temporal clusters (right).

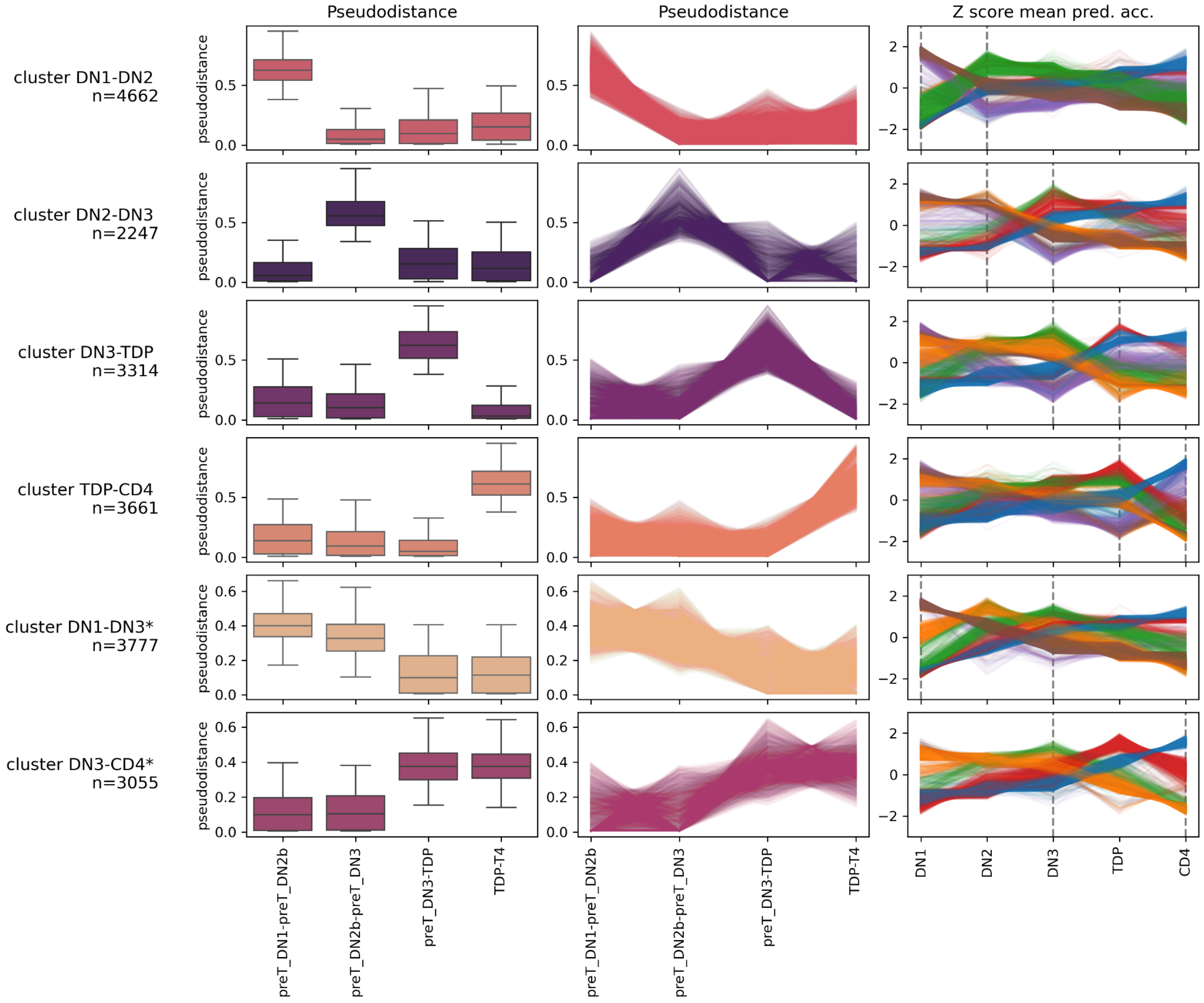
